# DLL3 expression predicts response and long-term outcome in small cell lung cancer patients treated with tarlatamab

**DOI:** 10.64898/2026.09.03.749238

**Authors:** Bingnan Zhang, Lingzhi Hong, Kaiwen Wang, C. Allison Stewart, Komal Shah, Wei-lien Wang, Alejandra Serrano, Luisa Solis Soto, Robert J. Cardnell, Alexa Halliday, Mitchell Parma, Elyse R. Lopez, Mukulika Bose, Cole Ruoff, Shafqat F. Ehsan, Kavya Ramkumar, Tim Li, Rosy Zheng, Loukia G. Karacosta, Jianjun Zhang, John V. Heymach, Carl M. Gay, Lauren A. Byers

## Abstract

**Background:** Small cell lung cancer (SCLC) is an aggressive, high-grade neuroendocrine carcinoma (hgNEC) with poor prognosis. Tarlatamab, a DLL3-targeting T-cell engager was approved in 2024 for relapsed SCLC, with almost doubled overall survival benefit compared to chemotherapy. However, over half of patients do not respond to tarlatamab and others develop resistance within months. With multiple DLL3-targeting therapies in clinical trials, there are currently no validated predictive biomarkers to identify patients most likely to benefit from tarlatamab. In this study, we evaluate baseline DLL3 IHC level in patients as a predictive biomarker to tarlatamab clinical outcome.

**Methods:** We assembled a cohort of 138 patients within IRB-approved MD Anderson GEMINI database (PA13-0589) with DLL3 expression by CLIA-validated immunohistochemistry (IHC), and those treated with tarlatamab from 7/1/2024 to 3/30/2026. Only those with DLL3 IHC results and treated with tarlatamab were included in this predictive biomarker study. All DLL3 levels were reported as percentage cells with cytoplasmic or surface DLL3 expression, including 24 with additional intensity score and H-score calculation. demographic, clinical, and outcome data were collected. Systemic and intracranial response data were assessed by RECIST and mRANO BM criteria respectively. Longitudinal liquid biopsies were taken from a subset of patients for circulating tumor DNA (ctDNA) and (Cytometry by Time-of-Flight) CyTOF analyses.

**Results:** In the predictive biomarker cohort of 54 patients with SCLC treated with tarlatamab, Median DLL3% level in tarlatamab treated cohort was 75% (ranges 0-100%). Median follow-up time was 9.9 months (95% CI, 7.1 – 12.2), median time on tarlatamab treatment (ToT) was 5.8 months (95% CI, 2.9 – NA). At the data cut-off, 61% (33 out of 54) patients were alive. Kaplan-Meier curve stratified by DLL3 % of <=75% vs. >75 showed significant longer ToT and a trend toward longer median OS (mOS). In DLL3>75% compared to DLL3 <=75%: mToT was 10.1 months vs. 2.9 months (p=0.036), mOS: not reached vs. 9.5 months (p= 0.066). Among 41 patients who completed one cycle of tarlatamab treatment and were eligible for systemic (extra-cranial) response per RECIST criteria; we observed partial response (PR) in 51% (21) patients, stable disease (SD) in 22% (9) patients; and progressive disease (PD) in 27% (11) patients. The median DLL3 % in those with PR vs. PD were 80% vs. 40% (p=0.006); median DLL3% comparing PR and SD were 80% vs. 60% (p=0.051). Among 35 patients evaluable for intracranial response, best objective response prior to any brain radiation were 5 (14%) CR,11(31%) PR, 8 (23%) SD and 11 (31%) PD. 9 received brain radiation (8 SRS, 1 WBRT) during tarlatamab to achieve better intracranial control while systemic control is maintained. DLL3% does not appear to be correlated with intracranial response. Longitudinal ctDNA and CyTOF analyses showed persistent ctDNA positivity, as well as increasing *NEUROD1*-subtype cells in circulating tumor cells in post-tarlatamab progression samples. We additionally profiled DLL3 expression by IHC in 138 patients with SCLC and found heterogenous DLL3 expression, with Median DLL3% of 70% (ranges 0-100%) across different biopsy sites.

**Conclusions:** Our data suggests DLL3 expression in patients with SCLC is heterogeneous and not ubiquitous, with median DLL3 IHC of 70%. DLL3 IHC predicts objective response and long-term outcome with tarlatamab. In addition, dynamic monitoring of ctDNA level and CTCs by CyTOF are suggestive of development of tarlatamab resistance, however further studies with larger cohort of paired longitudinal samples are needed to validate the novel blood-based biomarkers. To our knowledge, this study is the first to validate DLL3 IHC as a predictive biomarker for tarlatamab in SCLC, which leads the way for optimizing treatment selection and combinatorial therapies for the subset of patients less likely to respond to tarlatamab.

## Introduction

Small cell lung cancer (SCLC) is an aggressive, high-grade neuroendocrine carcinoma (hgNEC) characterized by early metastases, chemotherapy resistance and overall poor prognosis.^1^ Tarlatamab, a DLL3-targeting T-cell engager, was approved in 2024 for relapsed SCLC. However, the 35% objective response rate and 5.3 months median progression free survival (PFS) with tarlatamab in the confirmatory phase 3 study means over half of patients do not respond to tarlatamab and others develop resistance within months.^2^ To date, predictive biomarkers to tarlatamab response remain elusive.

DLL3 is an inhibitory notch ligand expressed at some level in over 90% of SCLC and many other neuroendocrine neoplasms, however, there is considerable heterogeneity in the expression levels that was previously under-explored.^3^ DLL3 expression was dismissed as a biomarker in earlier studies of tarlatamab efficacy due to some activity observed in DLL3 low or DLL3 negative cases, however both in DeLLphi300 and in tarlatamab’s FDA profiling, the data suggested increased DLL3 expression trended with higher magnitude of clinical benefit.^4, 5^ In addition, several recent studies using patients’ longitudinal blood samples and circulating tumor cells suggested DLL3 downregulation and T cell exhaustion may predict resistance to tarlatamab^6, 7^ Another recent study showcased possible resistance to tarlatamab associated with SCLC transcriptional subtypes, either in the *POU2F3* molecular subtype of SCLC with low DLL3 expression or via subtype switching from *ASCL1* to *NEUROD1* subtype accompanied by DLL3 down-regulation.^8^ Despite emerging evidence, it remains unclear whether DLL3 level in patients with SCLC predict outcome with tarlatamab treatment.

With the multitude of DLL3-targeting therapies in clinical trials, there is currently no validated predictive biomarker to identify patients most likely to benefit from tarlatamab, and vice versa, those less likely to respond may need alternative or combinatorial approaches. We hypothesize that analogous to PD-L1 level correlating with response to PD1/PD-L1 immune checkpoint blockade, there is inter-tumoral heterogeneity in DLL3 expression, and that baseline DLL3 IHC level can be used as a predictive biomarker for tarlatamab outcome.

## Method

### Study Population

The project was performed under The University of Texas, MD Anderson Cancer Center Institutional Review Board approved protocols PA13-0589 (GEMINI), PA14-0276 and PA16-0661 for clinical data, tissue and blood sample collections. From 3/2024 to 3/2026, 138 patients with ES-SCLC had tumor specimens tested for DLL3 by immunohistochemistry assays. Patients received tarlatamab who met the following criteria were included for predictive biomarker analysis: (1) diagnosis of pathologically confirmed ES-SCLC; (2) received at least one dose of tarlatamab. Patients with transformed or mixed SCLC were excluded. Clinical characteristics, pathology reports, imaging and outcomes were collected through medical records review. Longitudinal liquid biopsies were taken from a subset of patients for circulating tumor DNA (ctDNA) and (Cytometry by Time-of-Flight) CyTOF analyses.

### DLL3 expression staining

DLL3 IHC expression was assessed on tumor cells via CLIA-validated assay using Roche Ventana platform (anti-DLL3 antibody SP347) at MD Anderson. DLL3 expression was quantified based on the percentage of positive tumor cells (range, 0% - 100%). For a subset of patients, H-score quantification was performed by an expert pathologist (A.S) on the available DLL3 IHC slide, multiplying staining intensity (weak: 1+, moderate: 2+, strong: 3+) with percentage cells for each intensity to generate an H-score (H-score range, 0 - 300).

### Statistical analysis

When analyzed as a continuous variable, DLL3 was described by the median and interquartile range. Group comparisons were performed using the Mann-Whitney U (two groups) or Kruskal-Wallis tests (>= 3 groups). When analyzed by category with different cut-off thresholds, DLL3 was summarized by frequency and proportion. All hypotheses were tested in a two-sided fashion, and p less than 0.05 was considered statistically significant. RECIST version 1.1 was used to assess the systemic objective response. Time on treatment (ToT) was defined as the interval from the date of the first dose of tarlatamab to the date of the last dose. If a patient is still receiving tarlatamab at the time of data cutoff, the ToT is censored at the last known treatment date. overall survival (OS) was defined as the interval from the date of the first dose of tarlatamab to the date of death from any cause. Patients alive at last follow-up were censored for the OS analysis. The Kaplan-Meier method was used to estimate ToT and OS. Differences between groups were assessed through the log-rank test. Univariate and multivariate Cox proportional hazards (PHs) models were applied to further evaluate the association of covariates with survival outcomes.

### Circulating tumor DNA (ctDNA) and (Cytometry by Time-of-Flight) CyTOF analyses

Blood was collected into Streck tubes, plasma was isolated within 24 hours and stored at −80C in 2ml aliquots. Pre-tarlatamab and post-tarlatamab progression samples were identified, DNA was extracted from 2ml of plasma, and DNA was quantified using a Qubit 2.0 DNA HS Assay (Life Technologies, Grand Island, NY). PBMC isolation from whole blood, antibody panel design, CyTOF processing, CTC with SCLC subtype marker expression detection, and computational analysis for SCLC subtype assignment were performed as described previously.^9^ CTC subtype proportions were calculated for each patient sample as the percentage of cells assigned to each subtype relative to the total number of CTCs detected.

## Results

### Tarlatamab cohort characteristics and DLL3 level

54 patients with relapsed SCLC who received tarlatamab were included in the predictive analyses of DLL3 IHC level and outcome. In this cohort, median age was 65, and 56% were male. 92% had current or former tobacco exposure. 70% of patients received tarlatamab as second line treatment. 46% had Liver metastases and 78% had brain metastases present prior to tarlatamab administration. Notably, 61% had new brain metastatic lesions detected prior to tarlatamab initiation **(Table 1)**. DLL3 level in the tarlatamab cohort ranged from 0-100% by positive percent cells, with a median of 75%. Among those, 24 patients had additional DLL3 H-score analyses by expert pathologist review. Median H-score was 90, with ranges 0-285 **(Suppl. Fig.1A).**

**Figure 1:**
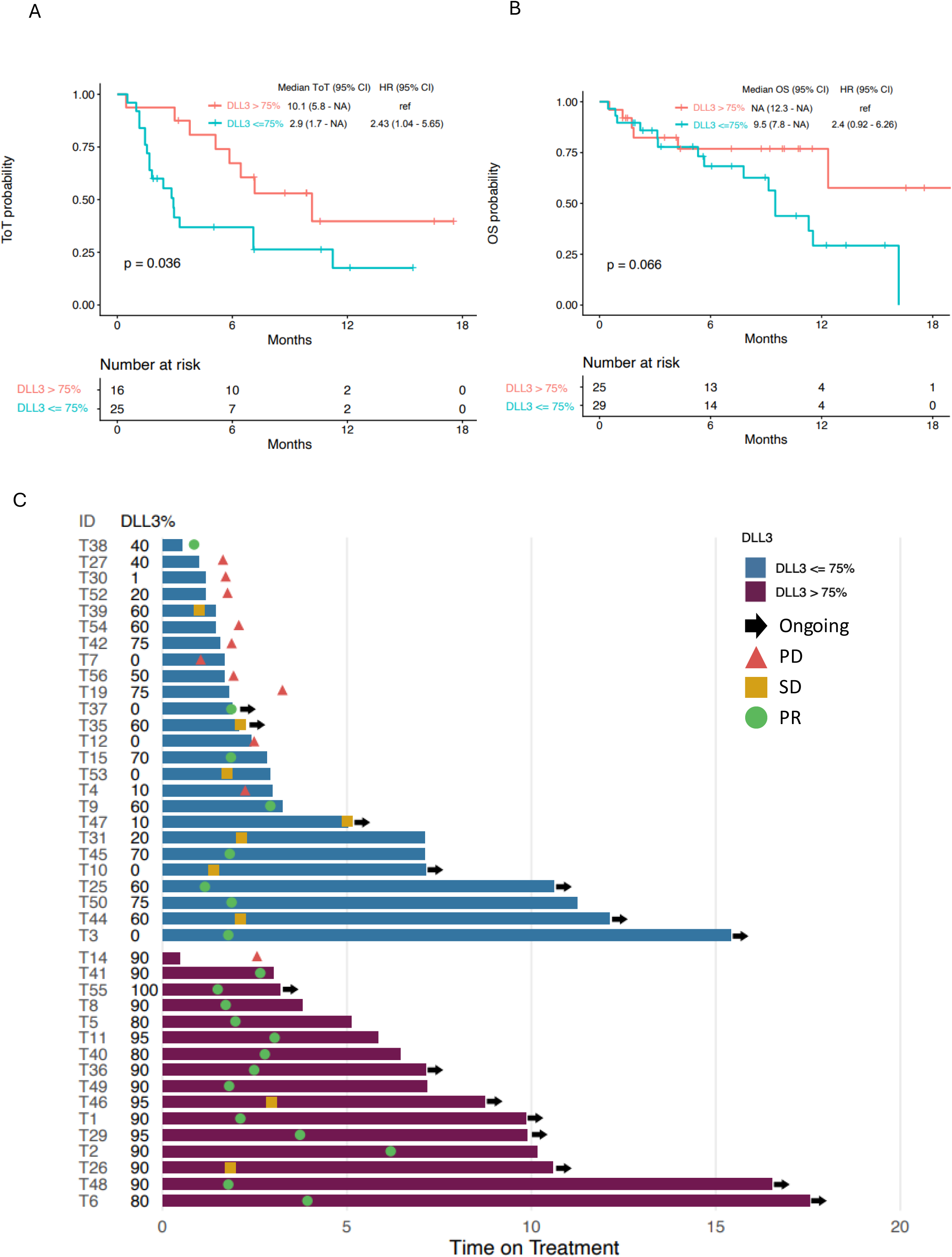
Outcomes in patients with extensive-stage small cell lung cancer treated with tarlatamab stratified by DLL3 expression at a cutoff of 75%. Kaplan-meier plot of A) time on treatment (ToT) and B) overall survival (OS). C) Swimmer plot of ToT. PD, progression disease; SD, stable disease; PR, partial response.

**Table 1.** Patient baseline characteristics.

| Parameters | N = 54 |
| --- | --- |
| Age, median years (range) | 65 (42 – 84) |
| Gender, n (%) |  |
| Male | 30 (56) |
| Female | 24 (44) |
| Initial Diagnosis | 46 (10-126) |
| ES-SCLC, n (%) | 46 (85) |
| LS-SCLC, n (%) | 8 (15) |
| Smoking, Median pack-year (range) | 46 (10-126) |
| Former/current, n (%) | 50 (92) |
| Never, n (%) | 4 (8) |
| DLL3 positive %, Median (range) | 75 (0-100) |
| 0% | 8 (15) |
| 1 – 50% | 9 (17) |
| 51 – 75% | 12 (22) |
| 76 – 100% | 25 (46) |
| Line of Tarlatamab, n (%) |  |
| 1* | 2 (4) |
| 2 | 38 (70) |
| 3+ | 14 (26) |
| Liver metastases, n (%) |  |
| Yes | 25 (46) |
| No | 29 (54) |
| Brain metastases (BM), n (%) |  |
| Yes | 42 (78) |
| New BM detected prior to tarlatamab | 33 (61) |
| No | 12 (22) |

\* Received tarlatamab during first line maintenance as part of a clinical trial.

### DLL3 level predicts long-term outcome on tarlatamab

At the data cut-off with a median follow-up time of 9.9 months (95% CI, 7.1 – 12.2), 61% (33) patients were alive. As a surrogate for real-world median PFS, we calculated the median time on tarlatamab treatment (ToT) for patients who completed at least one cycle of tarlatamab. Median ToT was 5.8 months (95% CI, 2.9 – NA), Kaplan-Meier curve stratified by DLL3% of 75% (median DLL3 of this cohort) showed significantly longer ToT with DLL3 >75%; 10.1 months (95% CI, 5.9 - NA) vs. only 2.9 months (95% CI, 1.7 - NA) in those with DLl3 <=75% (p=0.036) **(Fig.1A, 1C).** All patients who received at least one dose of tarlatamab were included in the survival analyses. In our cohort, a trend towards longer median OS was seen in patients with DLL3 >75%; mOS: not reached (95% CI, 12.3 - NA) vs. 9.5 months (95% CI, 7.8 - NA) (p= 0.066) **(Fig.1B).** Similarly, there was a trend towards longer ToT and longer OS using different DLL3 thresholds including 10%, 25% and 50%, with p-values approaching statistical significance as the thresholds become higher **(Suppl. Fig.2).** In univariable Cox analysis, higher DLL3 expression was associated with a trend toward improved overall survival (HR 0.92 per 10% increase, 95% CI 0.82 - 1.04; p = 0.19). In the 24 patients with H-score quantification, H-score of over 150 is associated with significantly better survival (12,3 mo vs. 5.5 mo, p=0.038) **(Suppl. Fig.1B)**. These data suggest higher DLL3 levels were associated with durability of treatment and better long-term survival in this cohort.

**Figure 2:**
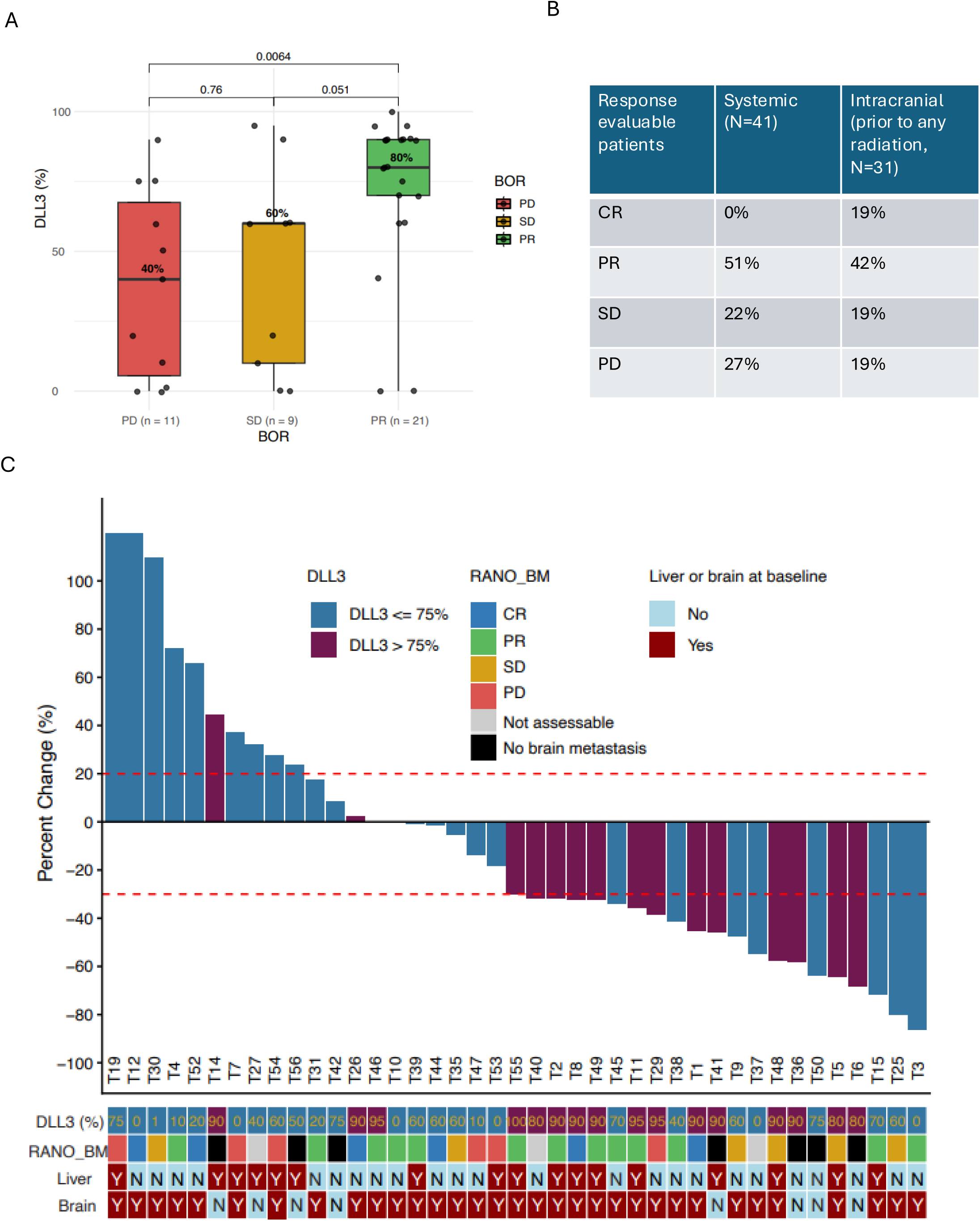
Association between DLL3 expression and best overall response (BOR), including systemic and intracranial responses, in patients with ES-SCLC treated with tarlatamab. A) Dot plot showing the association between DLL3 expression and systemic BOR. B) Systemic and intracranial BOR of the overall cohort. C) Waterfall plot showing the best percent change from baseline in extracranial target lesions.

### DLL3 level predicts systemic response to tarlatamab

41 patients completed at least one cycle of tarlatamab treatment and were assessed for systemic response using RECIST criteria. We observed partial response (PR) in 51% (21) patients, SD in 22% (9) patients; and PD 27% (11) patients. The median DLL3% in those achieving PR compared to PD were 80% vs. 40% (p=0.006); median DLL3% in PR compared SD were 80% vs. 60% (p=0.051) **(Fig.2A-B)**.

In addition, an expert neuro-radiologist graded intracranial response with tarlatamab using mRANO-BM criteria **(Fig.2B, 2C)**. In our cohort, 33 patients had new brain metastatic lesions prior to tarlatamab initiation. Among 31 patients evaluable for intracranial response, best objective responses to tarlatamab *prior to any* brain radiation were 19% (6) CR, 42% (13) PR, 19% (6) SD and 19% (6) PD. 29% (9) of patients received brain radiation during tarlatamab treatment to achieve better intracranial control, while ongoing tarlatamab continued to control extra-cranial disease. Systemic and intracranial responses were concordant in 77% (24/31) of cases. DLL3% by IHC does not appear to be correlated with intracranial response **(Fig.2C)**.

We also conducted univariant analyses of additional potential factors associated with tarlatamab benefit, incorporating clinical factors and baseline laboratory values within 24 hours prior to tarlatamab start. Baseline liver metastases was an unfavorable risk factor for tarlatamab in terms of ToT (Hazard ratio 2.61, 95%CI [1.13-6.02]), while absolute neutrophil count (ANC) greater than 3.6 x 10^3^, and high neutrophil to lymphocyte ratio (NLR<3.5), trended towards worse tarlatamab benefit for OS (**Suppl. Fig.3A,B**).

**Figure 3:**
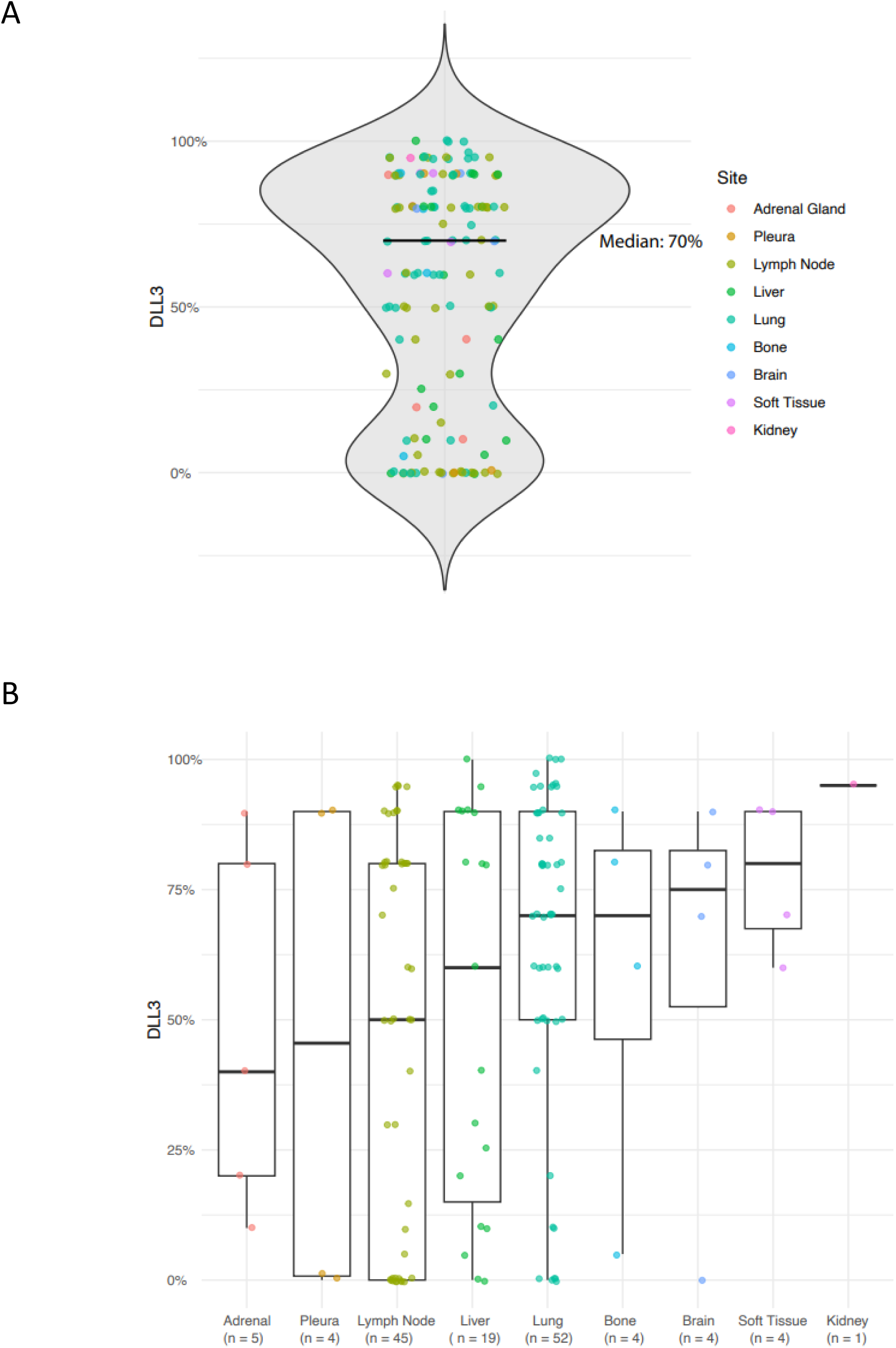
DLL3 expression by biopsy site in ES-SCLC (N = 138). A) Violin plot showing overall distribution. B) Box plot showing the distribution, with individual dots representing individual samples. The center line indicates the median, and the box represents the IQR.

Other clinical factors, such as whether tarlatamab was given as first or second line (line of tarlatamab <=2) versus later line (line of tarlatamab >2), or time interval between tarlatamab and last chemotherapy (TTC) more or less than 3 months did not show significant impact on ToT or OS **(Suppl. Fig.4A-D).**

### DLL3 IHC profiling in SCLC by biopsy site

In 138 patients with ES-SCLC tumor specimens tested for DLL3 our CLIA-certified IHC assay, DLL3 percentage cell positivity was heterogeneous among patients, ranging from 0 to 100%, with a median of 70% **(Fig. 4A)**. DLL3 level stratified by biopsy site showed majority of the biopsies were taken from lymph node, liver or lung, with numerically higher median DLL3 level in lung tissue samples **(Fig. 4B)**. However, statistical comparison of DLL3 levels by biopsy sites was limited due to relatively small numbers in several biopsy locations.

### CtDNA and CyTOF analyses suggest potential resistance mechanisms with SCLC molecular subtypes

Next, we analyzed blood samples from three patients in this cohort with either primary or acquired resistance to tarlatamab. Longitudinal samples from patients MDA-T11 (SC487, DLL3 95%), MDA-T50 (SC524, DLL3 95%), MDA-T4 (SC501, DLL3 10%) were taken at different timepoints during their tarlatamab treatment **(Suppl. Fig.5A)**.

For patient MDA-T11 (acquired resistance), the first blood sample (SC487-1) was collected 4 months prior to tarlatamab when patient was receiving first line immunotherapy maintenance. The ctDNA content was high despite no clinical progression by imaging **(Suppl. Fig.5B)**. The patient subsequently progressed on immunotherapy with significant systemic disease burden and numerous brain metastases and tarlatamab was started. While there were initial robust partial responses to tarlatamab including in the brain, after 5 months on therapy, the patient had intracranial progression (SC487-5) with lower but persistent ctDNA level, followed by systemic progression 6 weeks later (SC487-6). CyTOF analyses of blood sample SC587-6 showed 25 CTCs in the sample with majority of cells belonging to the SCLC-N subtype (**Suppl. Fig.5C**).

Similarly, patient MDA-T50 (acquired resistance) had a long treatment on tarlatamab with initial partial response, followed by oligo-progression after 3 months (SC524-1) that was treated with targeted radiation. Blood sample SC524-2 was collected 9 months after SC524-1 when the patient had multifocal progression. The rise of ctDNA comparing at the later timepoint likely reflected higher disease burden associated with progression **(Suppl. Fig.5B)**. CyTOF analyses of SC524-1 showed 13 CTCs with majority expressing *NEUROD1* subtype **(Suppl. Fig.5C)**.

Patient MDA-T4 (primary resistance) had baseline DLL3 IHC expression of 10% and was primary refractory to tarlatamab. Blood sample SC501-4 was collected immediately prior to tarlatamab, SC501-6 was collected 2 months on tarlatamab when the patient had small brain metastases and received stereotactic radiation, and SC501-7 was collected 1 month later when patient was confirmed to have systemic progression (SC501-7). CtDNA comparing SC501-4 and SC501-7 showed expected persistent and higher ctDNA content in post-tarlatamab progression setting **(Suppl. Fig.5B)**. Longitudinal CyTOF analysis showed predominant “triple-negative” phenotype of CTCs, with small proportions of *ASCL1* and *NEUROD1* expressing cells at baseline, while the proportion of *NEUROD1*-expressing cells did increase to a higher percentage at the later timepoint (SC501-6) **(Suppl. Fig.5C)**.

## Discussion

Our large cohort of patients with DLL3 IHC (n=138) demonstrated overall high DLL3 levels in all patients with SCLC across different tissue biopsy sites, with a median DLL3 level of 70%. To our knowledge, this is the largest cohort reported for DLL3 IHC characterization in SCLC. In the tarlatamab treated patients (n=54), median DLL3 level was 75%. For the first time, we demonstrated that baseline DLL3 level in our cohort predicts systemic objective response and long-term outcome to tarlatamab. Notably, those patients who achieved PR compared to SD and PD had statistically significantly higher DLL3% of 80%, vs 60 % and 40% respectively. When using the median DLL3 75% as cut-off, median ToT, as a surrogate for real-world PFS, was significantly longer, 10.1 months vs. 2.9 months (p=0.036). There was also a strong trend towards longer OS in the DLL3-high group (12.3 months vs. 9.5 months, p=0066). The ToT and OS trends hold true across lower DLL3% thresholds (10%, 25% and 50%) albeit with less statistical significance, suggestive of incremental benefit as DLL3 level increases. Similarly, in a subset of patients, DLL3 H-score of over 150 is associated with significantly better survival (12,3 mo vs. 5.5 mo, p=0.038). These data demonstrated that higher baseline DLL3 level predicted better clinical outcome with tarlatamab treatment in patients with SCLC.

Our findings are reminiscent of the debate surrounding the utility of PDL1 expression as a predictor of benefit to immune checkpoint inhibitors (ICIs). It is true that some PDL1-low patients will benefit from ICI therapy, but PDL1-high patients are more likely to benefit. PD-L1 level clearly matters, as PD-L1 testing is standard of care for many tumor types using ICIs, including in non-small cell lung cancer. In addition, for PD-L1 high (>50% expression), pembrolizumab has cancer agnostic approval for its single agent use. Similarly, it is true that some patients with DLL3 low expression still can respond to tarlatamab (possibly due to undetected expression heterogeneity and immune microenvironment factors), it is erroneous to suggest that DLL3 level does not matter in this setting. Importantly, identifying the group of patients who are less likely to benefit from tarlatamab as single agent is valuable, as it could help formulate superior combination strategies, such as with an antibody drug conjugate targeting DLL3 or, especially, in conjunction with agents targeting other surface antigens more highly expressed in DLL3-low patients. Additionally, obrixtamig (another DLL3 T cell engager) data in extra-pulmonary neuroendocrine carcinomas, already provided a compelling argument that DLL3 expression (cut-off 50%) by IHC clearly predicted response to obrixtamig with an objective response rate of 40% in DLL3 high (>=50%) vs. 3% in low (<50%)^10^. However, these emerging data do not suggest that systemic response to tartalamab is dependent on DLL3 levels alone. Several patients with high DLL3 levels still experienced primary resistance or early disease progression, and vice versa. As our CyTOF data and a recent preprint by Vasseur *et al*^8^ suggest, the emergence of NEUROD1 subtype of SCLC cells may be a mechanism of acquired resistance to tarlatamab, and immune related factors are also likely influencing either primary or acquired resistance^6, 7^. In addition, we found ctDNA changes correlated with clinical progression and the level of ctDNA likely reflected tumor disease burden in our patients.

Our study additionally delineated intracranial response using mRANO-BM criteria prior to any brain-directed radiation to not confound the data. We found that brain and extra-cranial systemic response were concordant in 77% of cases, however DLL3 level did not appear to be associated with brain response. In clinical practice as in this cohort, when there is ongoing systemic benefit despite intracranial progression, patients were able to receive brain-directed radiotherapy while continuing tarlatamab treatment for systemic benefit.

This study has several limitations. First, we acknowledge the limitations inherent to the retrospective data design including lack of control group and unmeasured confounding variables. In addition, we could not accurately report PFS given that we evaluated extra-cranial and intracranial responses separately; therefore, ToT was used as a surrogate for PFS. However, we acknowledge that ToT is likely over-estimating PFS, given in real-world situations, patients frequently receive radiation treatment to isolated intracranial progression or oligo-progression while continuing tarlatamab beyond progression.

In conclusion, we reported a large cohort of patients with SCLC and DLL3 expression profiling by IHC, and a separate cohort of patients with clinical outcomes treated with tarlatamab. Our data demonstrated for the first time that higher DLL3 level by IHC predicted tarlatamab benefit. We also showed other potential clinical and liquid biopsy biomarkers predictive of tarlatamab outcome. However, more studies are needed to elucidate the heterogeneity and dynamic changes of DLL3, as well as its inter-dependence with SCLC molecular subtypes, in terms of its predictive value in response to tarlatamab and emerging DLL3 targeted therapies. Building on these findings, further biomarker-enriching and potential combinatorial strategies will be needed to improve the outcome for subset of patients less likely to respond to DLL3 T cell engager therapy.

## Acknowledgement

This work was supported by: NIH K12CA088084 (B. Zhang), The NIH/NCI CCSG P30-CA016672; NIH/NCI P50-CA070907 (Byers); NIH/CHI R01-CA299261 (Gay), NIH/NCI R01-CA207295 (Byers); NIH/NCI R50-CA243698 (Stewart); NIH/NCI U01-CA256780 (Byers); NIH/NCI U24-CA213274 (Byers); The Department of Defense LC210510 (Byers); the American Lung Association Pierre Massion Lung Cancer Discovery Award (Byers), Lung Cancer Research Foundation (B.Zhang), the LUNGevity Foundation 2020-02 (Gay); CPRIT RP210159 (Gay); NETRF (Gay); and Rexanna’s Foundation for Fighting Lung Cancer (Byers, Gay, B.Zhang). We would also especially like to thank A.R.K., L.W.Y., M.J.A., R.B.N., K.E.N., J.O., J.K.R., B.& B.N, C.K., P.C.B., S.S., S.R. and W.A.B. for their philanthropic support of these projects.

## Supplementary Figures

**Supplemental Fig.1:**
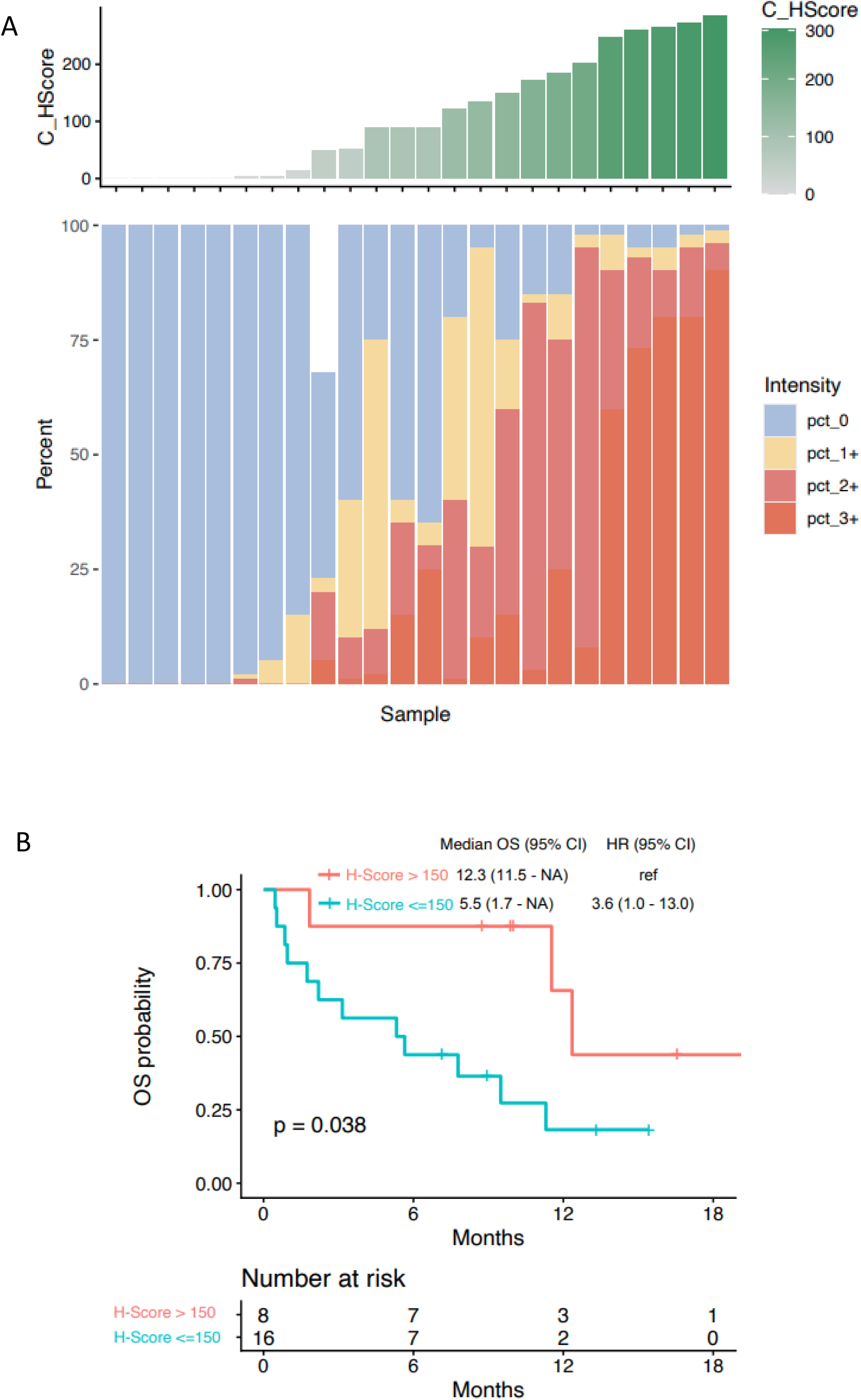
DLL3 immunohistochemistry (IHC) levels in ES-SCLC patients. A) DLL3 IHC cytoplasmic staining H-score and percentage of positive cells of 24 ES-SCLC patient tumor samples. B) Kaplan-meier plot of overall survival (OS) using a H-score cutoff of 90 (median value in the cohort) in patients. C). Kaplan-meier plot of OS using H-score cutoff of 150. HS: H-score

**Supplemental Fig.2:**
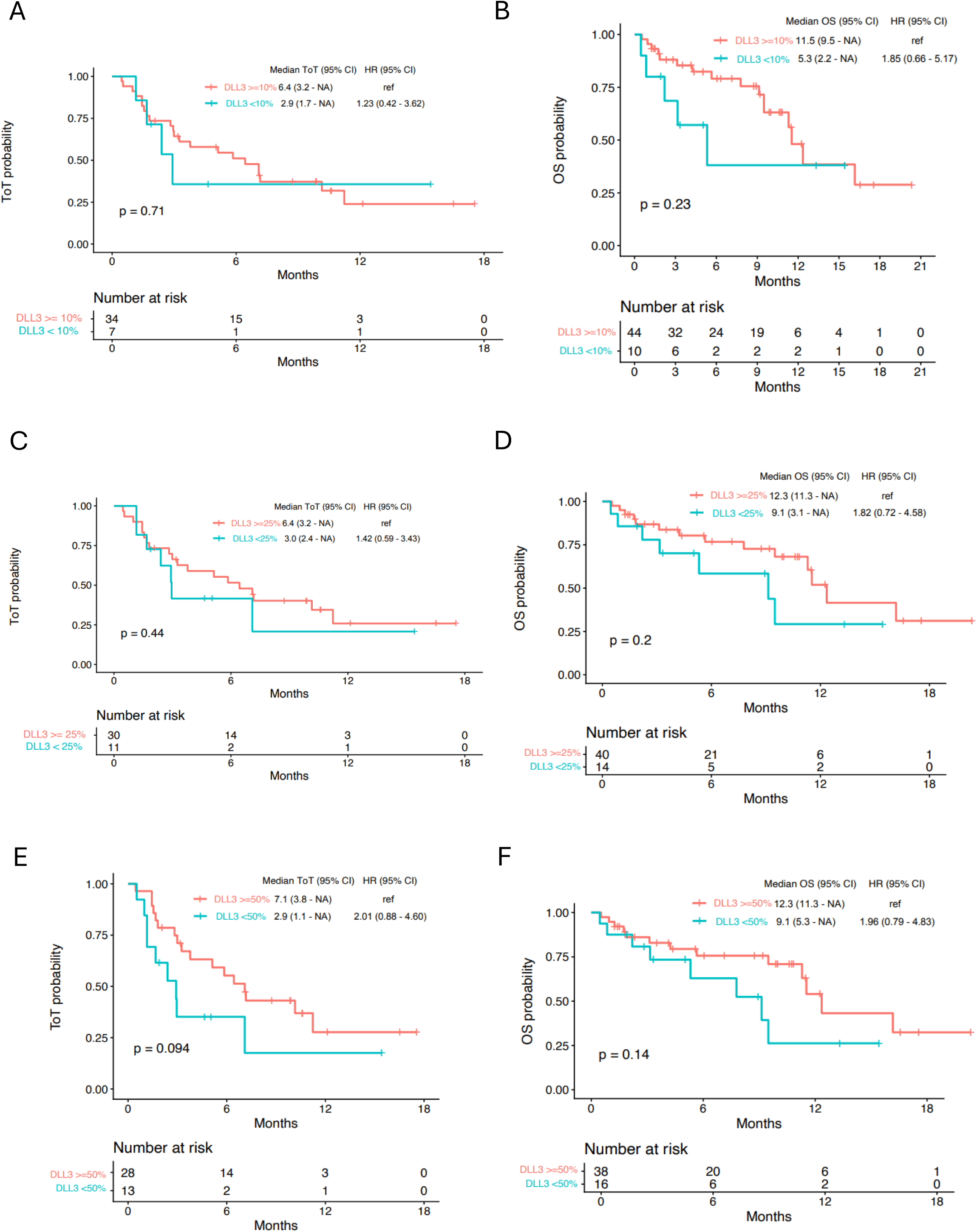
Outcomes in patients with ES-SCLC treated with tarlatamab stratified by different DLL3 expression cutoffs. Kaplan-meier plot of A) time on treatment (ToT) and B) overall survival (OS) with DLL3 at the cutoff of 10%; C) ToT and D) OS using a cutoff of 25%; E) ToT and F) OS using a cutoff of 50%.

**Supplemental Fig.3:**
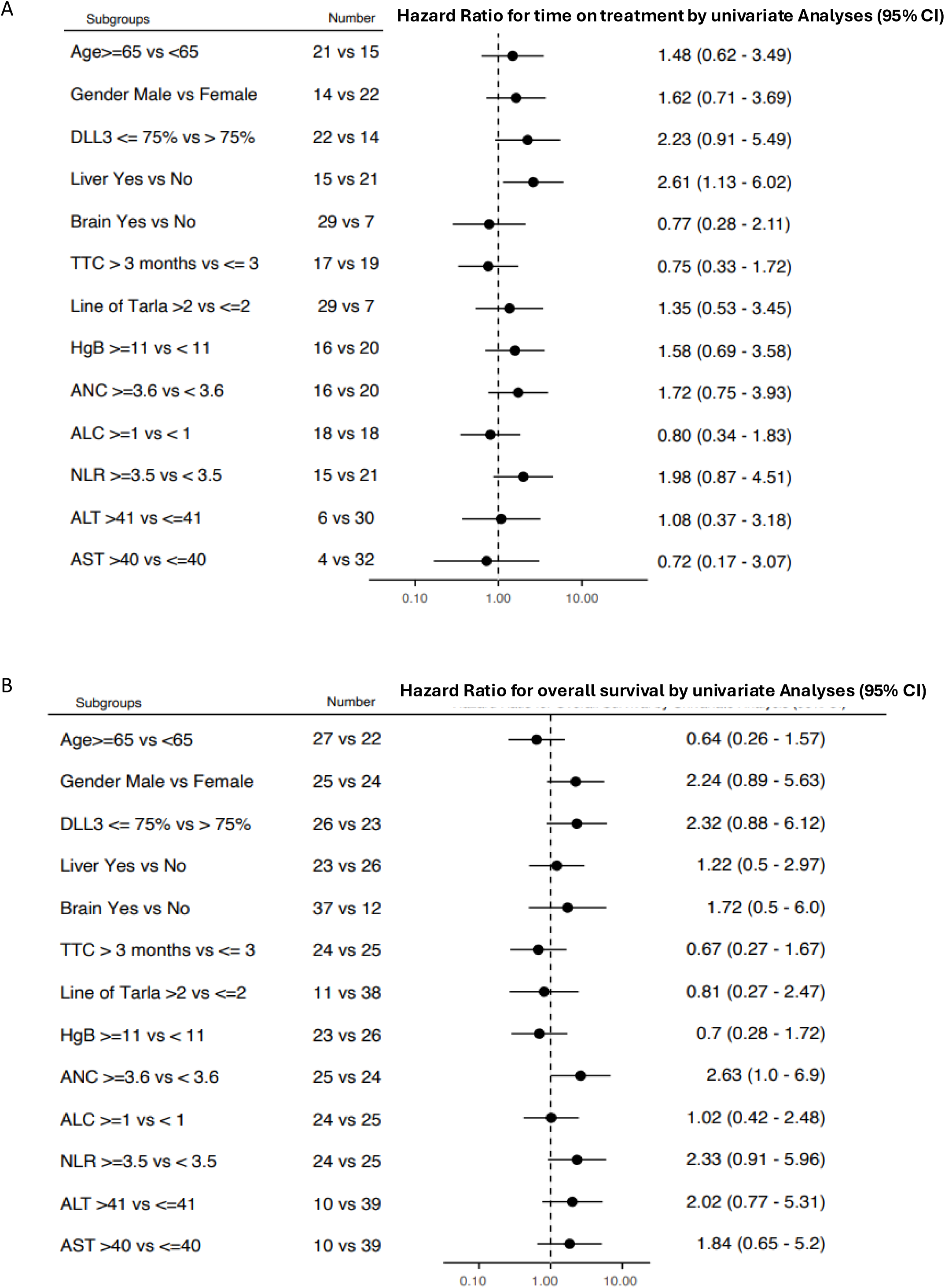
Clinical and laboratory predictors of outcome in patients with ES-SCLC treated with tarlatamab. Forest plot of variables and association using univariate analysis with A) ToT; B) Overall survival. ToT: Time on treatment, TTC: Time to last chemotherapy, Line of Tarlatamab>2: tarlatamab given after 2 lines of treatment, ANC: absolute neutrophil count (X10^3^/uL), ALC: absolute lymphocyte count (X10^3^/uL), NLR: neutrophil to lymphocyte ratio.

**Supplemental Fig. 4:**
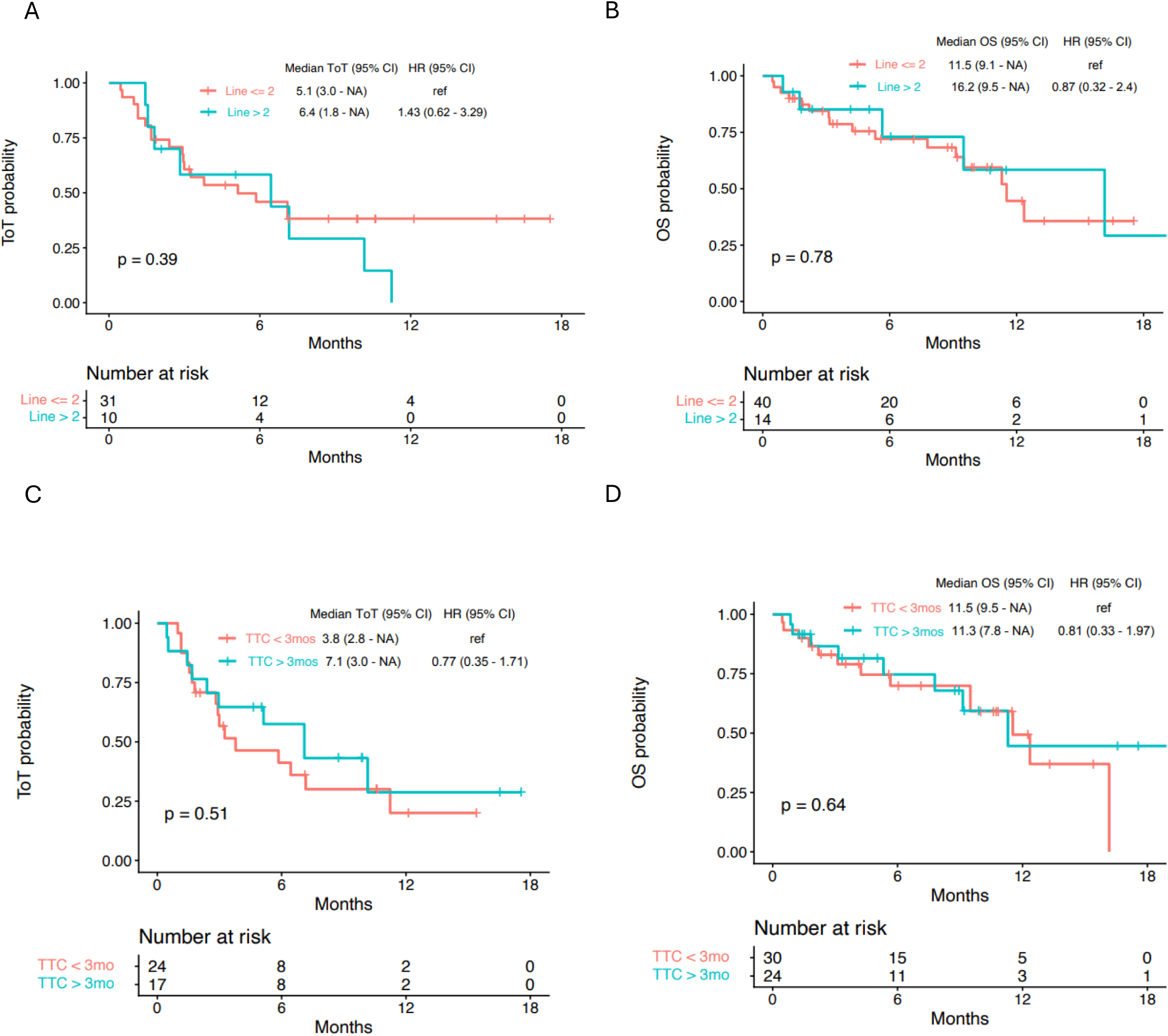
Outcomes in patients with ES-SCLC treated with tarlatamab stratified by line of tarlatamab and time to last chemotherapy (TTC). Kaplan-meier plot of A) time on treatment (ToT) and B) overall survival (OS) stratified by tarlatamab as first or second line, versus later than second line treatment; C) ToT and D) OS stratified by TTC.

**Supplemental Fig.5:**
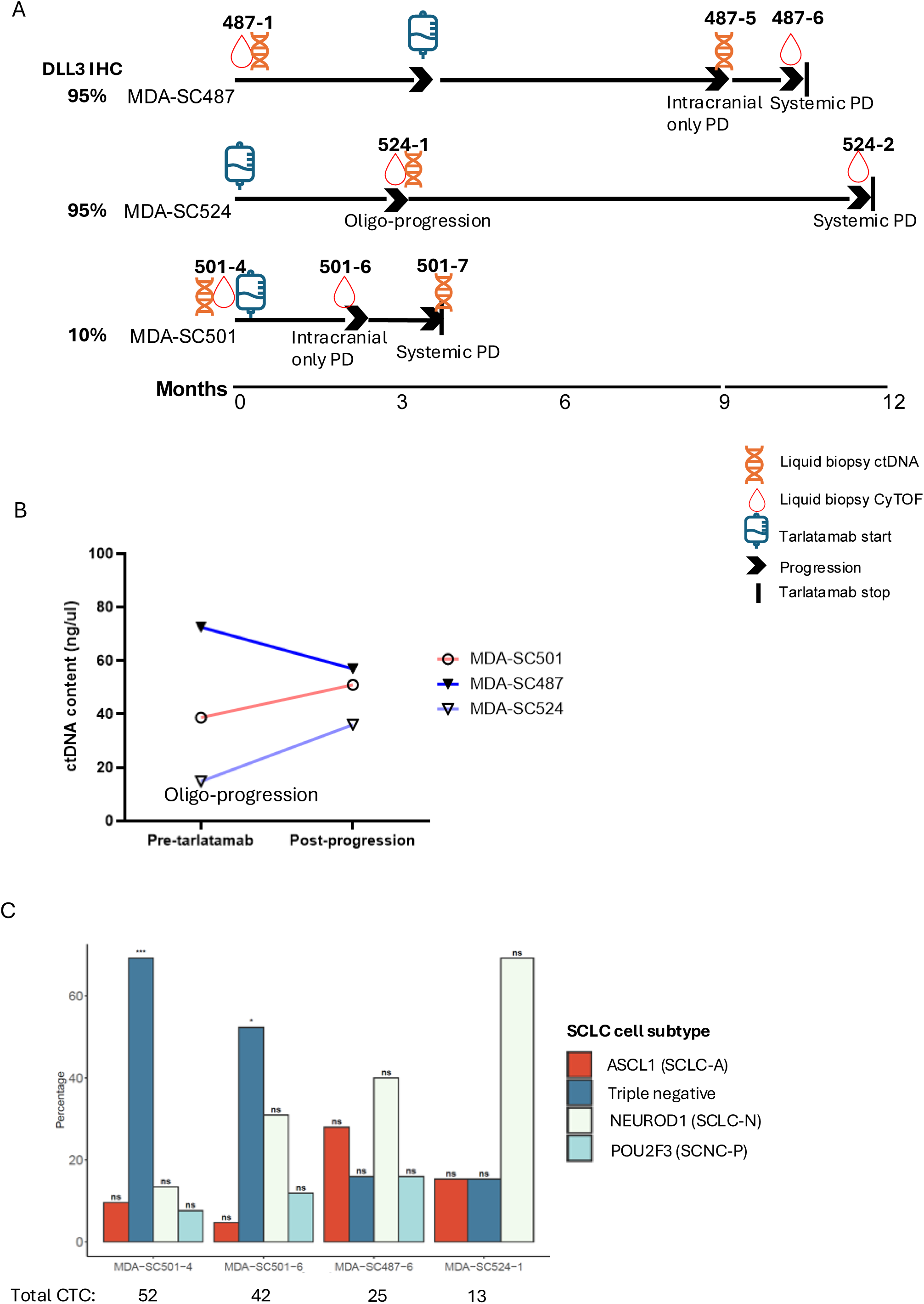
ctDNA and CYTOF analyses in patients treated with tarlatamab. A). Patient treatment journey, with baseline DLL3 IHC level, and sample collection timepoints. B). Changes in ctDNA content in the three paired patient samples on tarlatamab. C). Distribution of CTC molecular subtypes using CyTOF. A one-vs-rest Fisher’s exact test was performed for each subtype within each patient to determine whether the proportion of cells in a given subtype was significantly higher than that of the other subtypes. P-values were adjusted for multiple comparisons using the Benjamini-Hochberg False Discovery Rate method. Significance thresholds were defined as adjusted p-value < 0.05(*), < 0.01 (**), < 0.001 (***). C). ctDNA content in patient samples at various timepoints during tarlatamab treatment.

